# Adaptation to chronic malnutrition leads to reduced dependence on microbiota in *Drosophila*

**DOI:** 10.1101/113654

**Authors:** Berra Erkosar, Sylvain Kolly, Jan R. van der Meer, Tadeusz J. Kawecki

**Affiliations:** Department of Ecology and Evolution, University of Lausanne, CH-1015 Lausanne, Switzerland; Department of Fundamental Microbiology, University of Lausanne, CH-1015 Lausanne, Switzerland

**Keywords:** experimental evolution, microbiota, adaptation, juvenile development, digestion, nutritional stress, *Drosophila*

## Abstract

Numerous studies have shown that animal nutrition is tightly linked to gut microbiota, especially under nutritional stress. In *Drosophila*, microbiota are known to promote juvenile growth, development and survival on poor diets, mainly through enhanced digestion leading to changes in hormonal signaling. Here we show that this reliance on microbiota is greatly reduced in replicated *Drosophila* populations that adapted to a poor larval diet in the course of over 170 generations of experimental evolution. Protein and polysaccharide digestion in these malnutrition-adapted populations became much less dependent on colonization with microbiota. This was accompanied by changes in at least some targets of dFOXO transcription factor, which is a key regulator of cell growth and survival. Our study suggests that some metazoans have retained the evolutionary potential to adapt their physiology such that association with microbiota may become optional rather than essential.

## Introduction

Nutrient availability is a major factor limiting survival, growth and reproduction of many animal species ^1^, resulting in natural selection for adaptation to cope with nutritional stress. Yet, little is known about evolutionary adaptations that help juvenile animals not only to survive, but also grow, develop and reach maturity under chronic nutrient shortage. However, recent studies point to a particular importance of gut microbiota in coping with such chronic malnutrition. For example, mono-colonization with *Lactobacillus plantarum* buffers the growth of infant mice against the effects of nutrient shortage through a mechanism involving Insulin-like Growth Factor (IGF) signaling ^2^. Important insights about the mechanisms of microbiota-mediated enhancement of fitness under nutrient shortage have recently emerged from studies in *Drosophila.* As other insects that feed on a variety of sources, *Drosophila* have a rather simple and transient gut microbiota consisting of a subsample of ambient bacteria growing on their food (decomposing fruits)^3^. Nonetheless, as the more specialized commensals of mammals, these microbes provide a number of nutritional and metabolic benefits to their hosts ^4^,^5^. The same strain of *L. plantarum* that alleviated the effect of nutrient limitation on growth of mice ^2^ promotes the growth of *Drosophila* larvae on protein-poor diet, an effect mediated through upregulation of host's proteolytic enzymes leading to enhanced digestion and modulation of Insulin and TOR pathways ^6^,^7^. Another study using *Drosophila* also pointed to the commensal *Acetobacter pomorum* controlling larval growth by modulating Insulin/IGF-like signaling (IIS); this phenotype was again particularly pronounced on poor diets ^8^. Based on these findings, one might hypothesize that animal populations often exposed to chronic malnutrition would adapt by evolving an improved ability to benefit from their microbiota.

We address this hypothesis with experimental evolution ^9^. To study evolutionary adaptation to chronic juvenile malnutrition, we have maintained six outbred *Drosophila melanogaster* populations (“Selected” populations) for over 170 generations on an extremely poor larval diet (containing only 0.3% w/v of yeast). The nutrient content of the poor diet is so low that non-adapted larvae take twice as long to develop as on a standard diet and the resulting adults are only half the normal size ^10^. Compared to six “Control” populations maintained in parallel on a standard diet, the Selected populations evolved increased egg-to-adult survival, smaller critical size for metamorphosis initiation and faster development on the poor diet ^10,11^.

Here we test how this enhanced performance of the malnutrition-adapted Selected populations depends on interactions with microbiota and study the underlying physiological mechanisms. By manipulating the microbiota colonization status of larvae we demonstrate that, contrary to our expectations, these malnutrition-adapted populations became largely independent from microbiota for growth and survival on the poor diet. We show that protein and carbohydrate digestion in Selected larvae is much less affected by microbiota than in Controls, in spite of both types of larvae carrying microbiota of similar composition and abundance. Finally, our populations exhibit differential expression of some targets of the major cell growth regulator dFOXO ^12^. This indicates that site-specific function of dFOXO contributes to the physiological changes that result from adaptation to malnutrition, which compensates the microbiota effect in the non-adapted Control populations.

## Results

### Effect of microbiota on development and survival of experimentally evolved populations

We have maintained experimentally evolving Selected and Control populations on, respectively, a poor and a standard diet under a discrete-generation density-controlled regime for over 170 generations (i.e., over 10 years, see *Experimental Procedures*). This culture regime hindered vertical transmission of microbiota within populations from one generation to the next, and was conductive to exchange of microbes between populations as well as with the general environment of the climate room. Therefore, we did not expect one-to-one coevolution between the populations and their specific microbial communities. For this reason, to test the effects of microbiota we used a “common” microbiota inoculum collected from the feces of adults of all 12 populations (see *Experimental Procedures* for details). The “germ-free” (GF) flies were fed heat killed inoculum to control for potential effect of bacteria as food.

Larvae of our Selected populations had previously been reported to develop faster and survive better than Control larvae on poor diet (but not on standard diet) ^10,11^, a manifestation of their evolutionary adaptation to the poor diet; however, in those studies the colonization of the larvae by microbiota was not controlled and not assessed. We hypothesized that the improved performance of Selected larvae on the poor diet is at least in part mediated by an improved ability to benefit from interactions with microbiota. If so, one would predict that their superiority over Control larvae would diminish if they were deprived of the help of microbiota, i.e., in a GF state. To test this prediction, we compared the length of larval development and survival of Selected and Control populations in a GF state and when experimentally colonized with microbiota collected from adult feces. On the poor food, while Control larvae colonized with microbiota developed 40% faster and were three times more likely to survive than their GF siblings, the corresponding effect of microbiota treatment on Selected larvae was much smaller (**Fig 1A, B**). On the standard food, the effect of microbiota on development and survival was markedly smaller (**Fig S1A, B**). Thus, while in the GF state the Selected larvae took 30% less time to pupate on poor diet and were about three times as likely to survive as the Controls, the advantage of the Selected over Control populations for both traits diminished when the larvae were colonized with microbiota (**Fig 1A, B**). In particular, the developmental time became statistically indistinguishable between the Selected and Control larvae in the colonized state (even though Selected still tended to develop about one day faster than Controls). These results are opposite to our prediction. They imply that, in the course of their evolutionary adaptation to the poor diet, the Selected populations became less dependent on microbiota and much better able to cope with nutrient shortage without their help.

**Fig 1.**
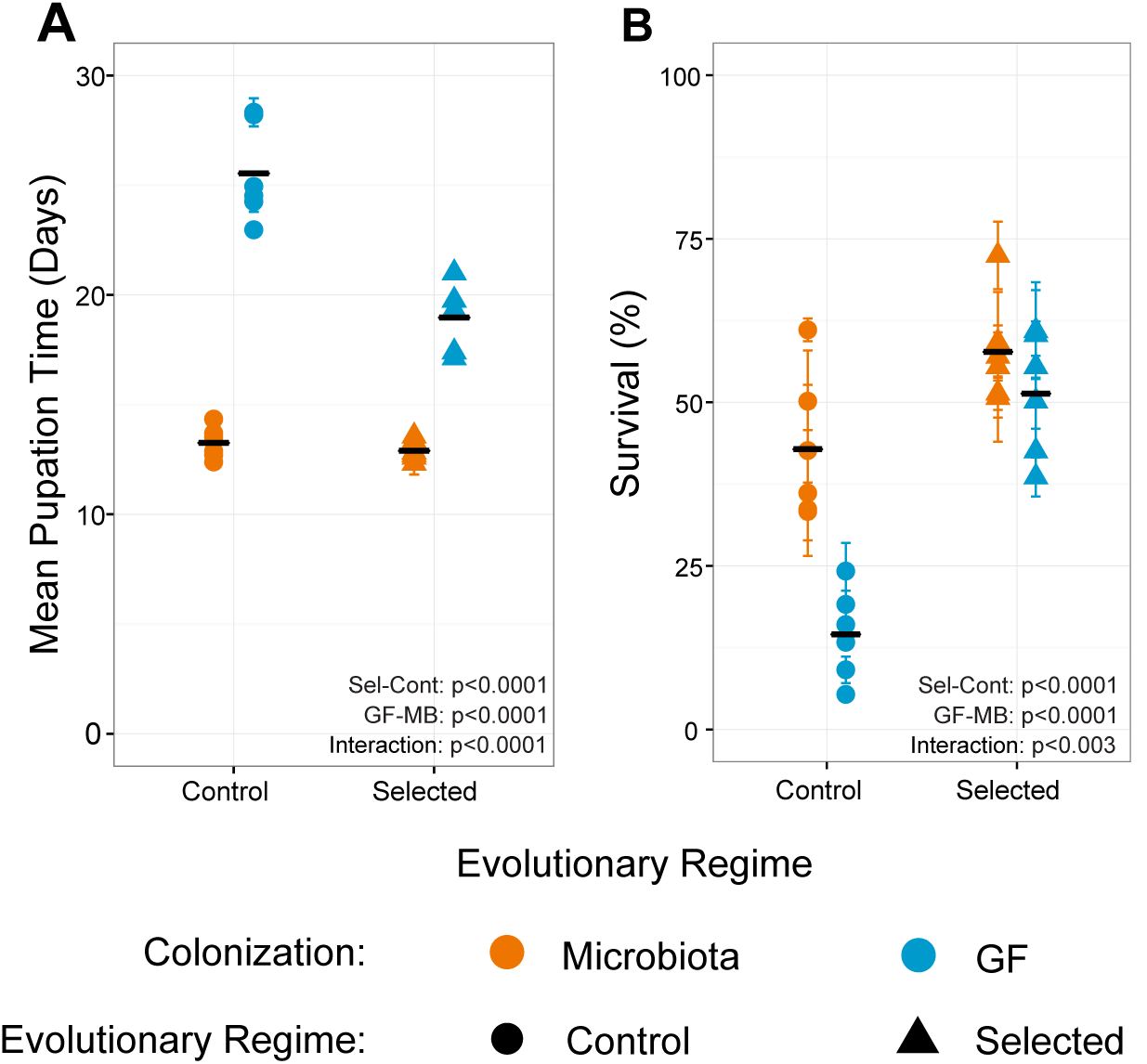
Microbiota affects development and survival differently in Selected and Control populations on poor food. **A.** Mean egg-to-pupa development time in Selected and Control populations, with or without microbiota. **B.** Mean egg-to-pupa survival rate under the same conditions. Symbols and error bars represent mean ± SEM for each population (where error bars are not visible, they are smaller than the symbols). Black horizonal bars represent the means of the six replicate populations. Main effect differences analyzed by GMM are represented in the panels. Interaction = Colonization × Regime. Detailed statistics are presented in Supplementary Table S1.

### Protein digestion in Selected and Control populations

Recently, it has been shown that upon nutrient scarcity, one of the members of *Drosophila* microbiota, *Lactobacillus plantarum*, promotes intestinal protease expression, leading to enhanced dietary protein digestion and increased amino acid concentrations in the host tissues ^7^. We therefore hypothesized that the weaker effect of microbiota on the survival and developmental time of Selected than Control populations could be mediated by a differential effect on protein digestion efficiency. To test this prediction, we measured protease activity (relative to the total protein content) in whole GF and microbiota-colonized larvae at different time points during the 3^rd^ instar (**Fig 2A**). This relative protease activity declined over time, which could reflect changes in protease secretion as well as an increase in the amount of protein accumulated in the larval body relative to the size of the digestive system. Irrespective of this apparent decline over time, inoculation with microbiota strongly enhanced proteolysis in Control larvae, but had a significantly smaller effect on proteolysis in Selected larvae. Thus, while GF Selected larvae exhibited (marginally significantly, *p* =0.083) higher levels of protease activity than GF Controls (particularly in mid- to late 3rd instar), the trend went in the opposite direction in microbiota-associated larvae. Thus, the pattern of proteolytic activity of the Selected and Control larvae in the absence and presence of microbiota matches the pattern of larval performance reported above. Apparently, being supplemented with microbiota helps Control populations to recover from low digestive activity, helping them grow and survive.

**Fig 2.**
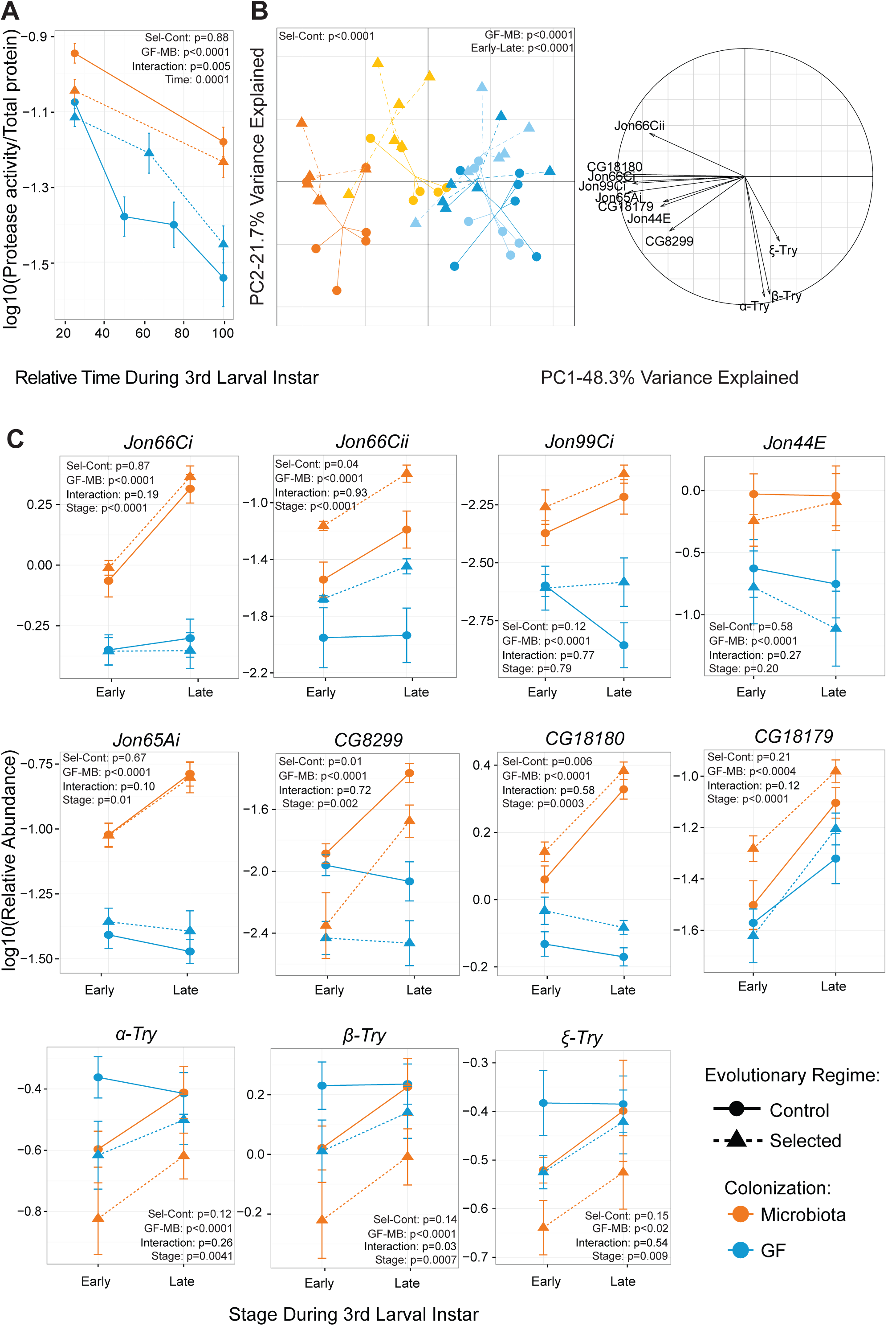
Microbiota affects protein digestion differently in Selected and Control populations. **A**. Protease activity in Selected and Control larvae through the 3^rd^ larval instar in the presence or absence of microbiota. **B.** Projections of protease expression dataset into 1^st^ and 2^nd^ PCs (left) together with correlation circle (right) representing the variables. <u>Light shade</u>: early 3^rd^ instar, <u>dark shade</u>, late 3^rd^ instar. **C.** Relative abundance (2^-ΔCt^) of different proteases measured by qRT-PCR from dissected guts of Selected and Control larvae at early and late L3 stage. Symbols represent mean ± SEM of the six replicate populations, with 3 biological replicates per population. A selection of key statistical results from GMM is represented in the panels. Interaction = Colonization × Regime. Detailed statistics including pairwise contrasts are presented in Supplementary Table S2.

As previously reported, seven serine proteases, including five Jonah proteases are transcriptionally induced upon colonization with *L. plantarum* ^7^. To determine if differences in expression of the same proteases may be responsible for the pattern of proteolytic activity in our populations, we dissected the intestines of GF and colonized larvae at early and late 3^rd^ instar, and carried out a qRT-PCR analysis on 11 proteases including Trypsins, Jonah proteases and a few others known to have serine type protease activity. Consistent with the previous report ^7^, we detected an elevated expression in all populations upon colonization with microbiota in all five Jonah proteases and three other serine proteases (*CG18179, CG18180, CG8299;* **Fig 2C**). However, trypsin superfamily proteases (*α-Try, β-Try, ε-Try*) (which are clustered together in the genome and reported to have a very localized expression in the gut ^13^) exhibited the opposite pattern, i.e., were downregulated by microbiota (**Fig 2C**). Out of the 11 proteases, we identified two (*Jon66Cii, CG18180*) whose mRNA levels were consistently higher in Selected populations compared to Controls; we also observed that *CG8299* had higher expression in Control than selected populations (**Fig 2C**). Trends for differences between Selected and Control populations could also be observed for several other proteases (all three Trypsins, *CG18179, Jon65Ai, Jon44E, Jon99Ci*, **Fig 2C**), but they were not sufficiently consistent between time points or replicate populations to be statistically significant. The digestive proteases are likely to some degree functionally redundant, and thus it is conceivable that evolution would achieve functionally similar changes in digestion by targeting different genes in different replicate populations, making detection of a signature of evolution in a gene-by-gene analysis difficult. Therefore, we analyzed the entire protease expression dataset with multivariate analysis of variance (MANOVA) and Principal Component Analysis (PCA). The correlation circle clearly confirmed that the levels of expression of the three Trypsins were positively correlated and well separated from other proteases (**Fig 2B right**). This suggests that these two groups of proteases are regulated by different processes and/or may have a different function within the gut. GF and microbiota-colonized larvae were clearly separated by the 1^st^ PC, with Selected and Control populations somewhat less distinctly separated along the 2^nd^ PC (**Fig 2B left**). Given that the 1^st^ PC explains more than twice as much variance as the 2^nd^, this implies microbiota are a major factor that changes protease levels, with a greater impact than the evolutionary history. Importantly, in spite of highly significant main effects of both microbiota and evolutionary regime in the MANOVA, there was no interaction between them (**Table S2**). Thus, even though these results suggest that evolutionary adaptation to poor food was in part mediated by changes in protein digestion, changes in the expression of digestive proteases cannot fully account for the differential effects of microbiota on the developmental time and survival.

### Carbohydrate digestion in Selected and Control populations

Given that our poor diet is low in carbohydrate as well as protein content, we next asked if carbohydrate digestion is also different between Selected and Control populations and if it is differentially influenced by microbiota. About 30 % of carbohydrates in both poor and standard diet consist of polysaccharides (starch) from the cornmeal (the rest are sucrose and glucose). Polysaccharide digestion occurs as a two-step process whereby starches are first broken-down to disaccharides by amylases before being hydrolyzed to monosaccharides. Alpha-amylase activity is under direct negative regulation by glucose concentration in *Drosophila* larvae, which occurs at the transcriptional level. Amylase activity is therefore expected to be lower in larvae with higher glucose concentration ^14,15^. We quantified amylase activity rates in Selected and Control larvae in both colonized and GF states. Microbiota had a striking effect on how amylase activity (again normalized to total larval protein content) changed over time: while it declined between an early and late 3rd stage in the microbiota-colonized larvae, it increased sharply during the corresponding developmental period in GF larvae (slope difference p < 0001, **Fig 3A**). Because no such increase is observed for protease activity (**Fig 2A**), it implies that GF larvae upregulate their investment in polysaccharide digestion relative to protein digestion towards the end of their development. Irrespective of these temporal changes, GF Selected larvae consistently showed three-fold lower amylase activity than GF Control larvae of the same stage (blue symbols in **Fig 3A**); this difference is much smaller and non-significant in microbiota-colonized larvae (orange symbols in **Fig 3A**). Thus, we again observed a pattern of interaction such that the difference due to evolutionary history was more pronounced in germ free than in microbiota-colonized state. However, fast development and high survival on poor diet (**Fig 1**) were associated with lower amylase activity. This implies that increased amylase activity is a sign of nutritional stress. Given the negative regulation of amylase activity by glucose concentration ^14,15^, these results suggest that Control larvae may have lower glucose levels than Selected larvae under GF conditions.

**Fig 3.**
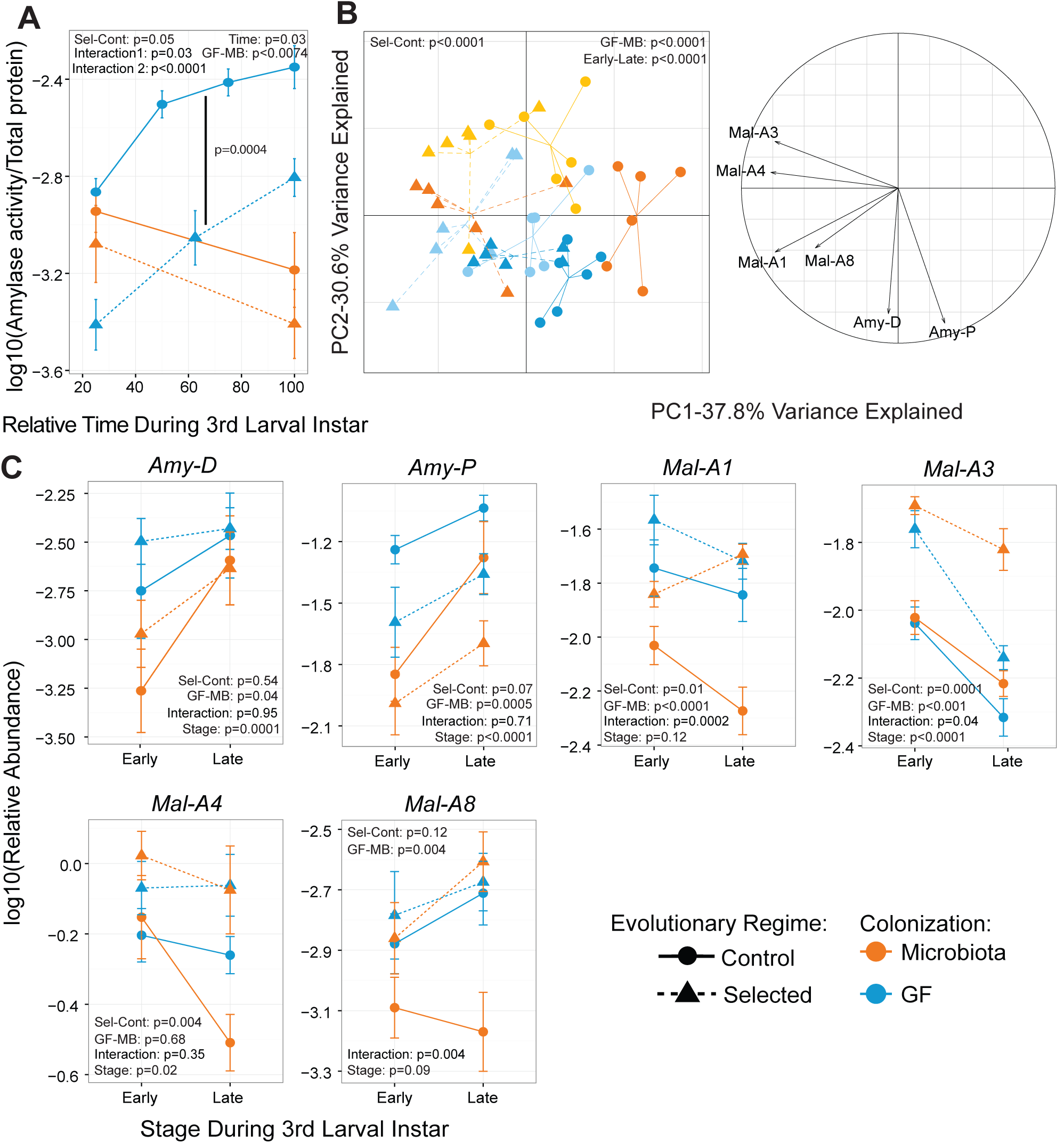
Microbiota affects carbohydrate digestion differently in Selected and Control populations. **A**. Amylase activity in Selected and Control larvae through the 3^rd^ larval instar in the presence or absence of microbiota. Significant pairwise difference between GF Control and GF Selected populations are shown with a black line. **B.** Projections of amylase and maltase expression dataset into 1^st^ and 2^nd^ PCs (left) together with correlation circle (right) representing the variables. <u>Light shade</u>: early 3^rd^ instar, <u>dark shade</u>: late 3^rd^ instar. **C.** Relative abundance (2^-ΔCt^) of different amylases and maltases measured by qRT-PCR from dissected guts of Selected and Control larvae at early and late 3^rd^ instar. Symbols represent mean ± SEM of for the six replicate populations, with three biological replicates each. A selection of A selection of key statistical results from GMM is represented in the panels. Interaction = Colonization × Regime, Interaction 2 = Time × Colonization. Detailed statistics including pairwise contrasts are presented in Supplementary Table S3.

To verify if the pattern we observed is regulated at the transcriptional level we quantified amylase transcript levels in the guts. We analyzed expression of two amylases. Both gene transcripts were significantly reduced in colonized larvae in all populations (**Fig3 C**). Under GF condition, *Amy-P* levels were higher in Control populations than in Selected populations, but no significant difference was detected in *Amy-D* levels (**Fig 3C**). Given that relative expression abundance of *Amy-P* is much higher than *Amy-D* (roughly 20 times, **Fig 3C**), *Amy-P* is likely to be the major gene contributing to the amylase activity pattern that we observed earlier (**Fig 3A**). Even though *Amy-P* expression is reduced by microbiota and is expressed at lower levels in Selected than Control populations, the expression pattern does not fully explain what we observe for amylase activity, and other regulatory mechanisms (e.g. cAMP levels ^14^) may also play a role in regulating amylase activity.

In the gut, glucose is generated through the hydrolysis of maltoses by maltases. If amylase activity is lower in Selected populations and upon microbiota colonization because of glucose concentration in the gut and/or hemolymph, maltase activity is predicted to be higher in these conditions. To check this we also analyzed expression of four *maltase* genes. In agreement with this prediction, we observed a high expression of maltases in Selected populations for *Mal-A1, -A3 and -A4*, although not for *Mal-A8* (**Fig 3C**). A consistent decrease in expression can be observed upon colonization only in Control populations for *Mal-A1, -A8* (**Fig 3C**). *Mal-A4* exhibits this trend only at late 3^rd^ instar but this is not statistically significant due to high variation among populations (**Fig 3C**). *Mal-A3* expression is rather induced in Selected populations upon colonization, and remains unchanged in the Control ones (**Fig 3C**). To spot the general trend among these carbohydrate-digesting enzymes we performed multivariate analyses. We observed a clear separation between the evolutionary regime, colonization status and developmental stage (**Fig 3B** left, **Table S3**). However, we observed only a marginally significant interaction between the evolutionary regime and developmental stage, and no interactions between other factors (**Fig 3B** left, **Table S3**). Furthermore, PCA correlation circle on carbohydrate digesting enzymes shows that amylase and maltase expression patterns are uncorrelated (**Fig 3B** right). Altogether, although Selected and Control populations exhibit different levels of carbohydrate-digesting enzymes, transcriptional differences of digestive enzymes cannot fully explain the interaction between evolutionary history and colonization status.

### Characterization of the microbiota

To identify the bacterial taxa that might be involved in the digestive enzyme induction in our experiments, we performed a microbial community profiling using 16S rRNA gene sequencing. All our populations carry the endosymbiont *Wolbachia* (data not shown), which would dominate the reads if the 16S sequencing were performed on DNA extracted from whole flies or from dissected guts. Therefore, we performed this analysis first using the feces of the adults pooled from the 12 populations, i.e., the microbial community used for experimental inoculations described above. This community was dominated by a single *Acetobacter sp.* that contributed ~82% reads, with ~14% reads attributable to Pseudomonadaceae, ~2% to other Acetobacteraceae and ~2% other less abundant taxa (**Fig4A, lowermost bar**).

**Fig 4.**
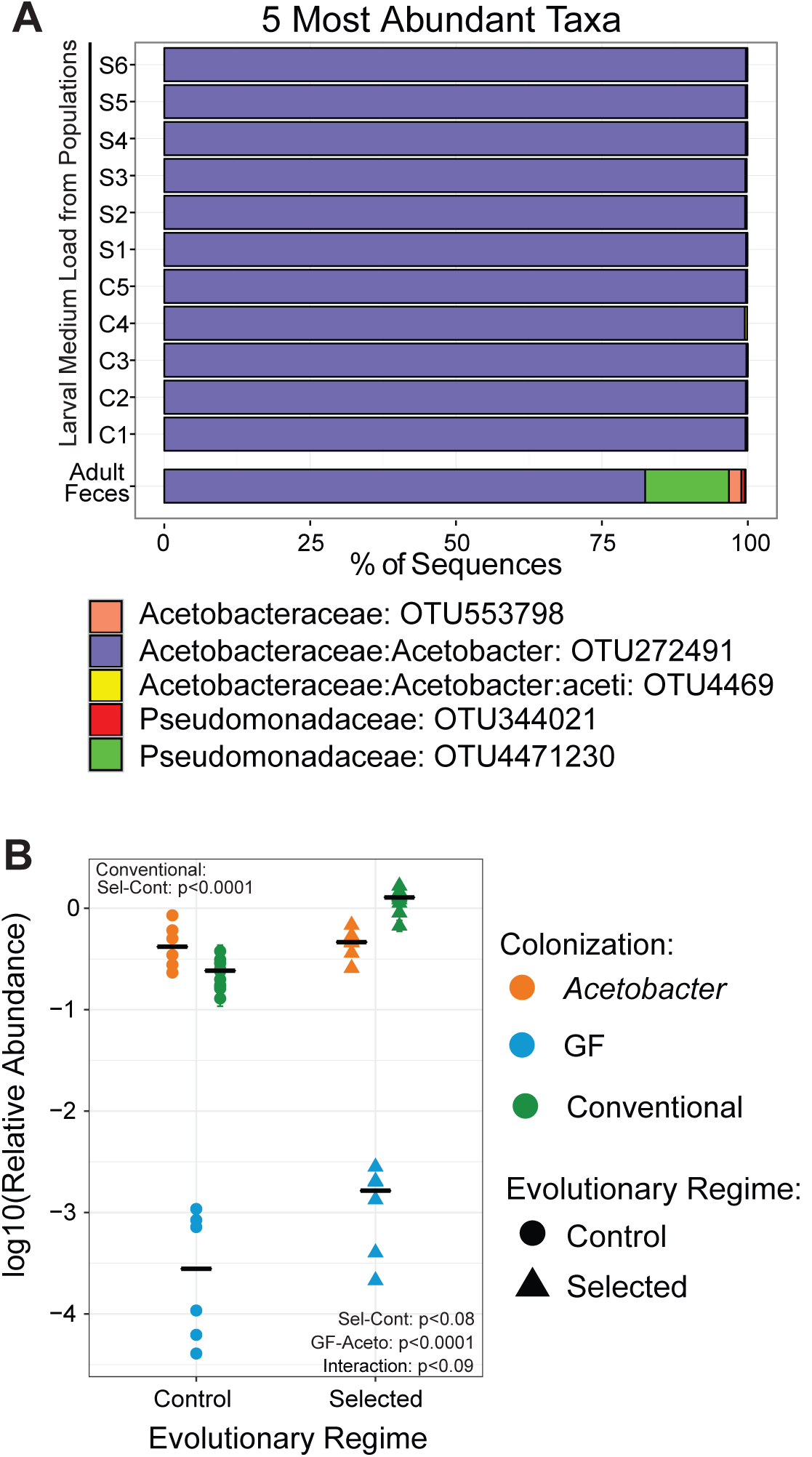
Microbiota of Selected and Control populations. **A.** The identities and relative abundances of 5 most abundant taxa in the mixed adult feces (used as the source of inoculum), and from the larval poor medium of Selected and Control populations previously colonized with that inoculum, assigned by 16S rRNA gene amplicon sequencing. C: Control, S: Selected Populations. **B.** The abundance of *Acetobaceraceae* relative to the host DNA in GF, *Acetobacter* mono-associated and conventionally reared (by experimental evolution) Selected and Control populations measured by qPCR. Symbols represent mean±SEM of for each population. Black bars represent the mean of the six populations within regime. Main effect differences analyzed by GMM are represented in the panel. Interaction refers to colonization × evolutionary regime. Details including pairwise contrasts are presented in Supplementary Table S4.

To check if association with Control versus Selected larvae promoted different members of this microbial community, we used this feces suspension to inoculate poor-diet larval cultures of each population upon hatching, and collected samples of the medium at the end of larval development (this was done in the same experiment that provided larvae for the gene expression experiment described above). 16S sequencing of these samples revealed that, irrespective of evolutionary history of the populations, they all consisted almost exclusively (> 99% of the community) of the single *Acetobacter* spp already prevalent in the inoculum (**Fig4A**).

Could then the differences between Selected and Control populations in the effects of microbiota inoculation on larval performance and digestive enzymes be mediated by differential colonization of their guts by this dominant *Acetobacter* strain? To address this question, we mono-colonized freshly hatched GF larvae of all twelve populations with this strain, allowed them to develop on the poor diet, and estimated the amount of bacteria inside the larval gut at the end of larval development. This was done by using qPCR to quantify bacterial DNA (using primers specific to Acetobacteraceae 16S rRNA gene) relative to host genomic DNA (using primers for *Actin*). We found no systematic difference between these experimentally colonized Selected and Control larvae in the amount of bacterial DNA relative to host DNA (**Fig 4B** orange symbols), nor in the absolute Ct values for the bacterial DNA (**Fig S1**). The latter indicates that the amount of bacterial DNA in these samples was about 1000-fold above the detection threshold; based on preliminary data (not shown) this roughly corresponds to 600-900 CFUs per larvae. Analogous Ct values for GF larvae were comparable to what was observed in a mock sample only containing sterilized water, which sets the detections limit (black line in **Fig S1**). This assures that our procedure of generating GF animals was effective.

The above results indicate that Selected and Control populations become similarly colonized by the dominant *Acetobacter* strain upon experimental inoculation followed by development on the poor diet. This implies that adaptation of Selected populations to poor diet did not cause any changes in the gut that would affect its colonization by commensals. However, this does not preclude a difference in the amount of bacteria they normally harbor under their respective evolutionary regimes (in their “conventional” environment), given that the regimes differ in diet and does not involve experimental inoculation. To address this issue, we used the same approach to quantify bacterial colonization by *Acetobacter* in the main cultures used to propagate these populations under the experimental evolution that is ongoing in the lab (i.e., on poor diet for Selected and on standard diet for Control populations). Interestingly, despite the difference in diet, these larvae reared in their respective conventional environments were colonized with comparable levels of Acetobacteraceae (**Fig 4B** green symbols). This suggests that the ability of Selected lines to become largely independent of microbiota (i.e. their ability to cope with being GF) is a physiological result of being adapted to malnutrition and not of being maintained GF by coincidence.

### Growth rate and activation of dFOXO targets

*Acetobacter pomorum* has been shown to promote larval growth through induction of Insulin/IGF-like signaling (IIS) by acetic acid secretion, evidenced by cytoplasmic retention of dFOXO in larval fat body ^8^. *Acetobacter sp*. in our system is also likely to secrete acetic acid since we observe a clear reduction from pH 3.5 to pH 2.0 in the media of all 12 populations upon colonization. We thus hypothesized that microbiota would promote larval growth in Control populations, but less so in Selected populations. However, adult size is thus not a good proxy for larval growth rate in these populations: because Selected populations evolved a smaller critical size for metamorphosis initiation, they reach a smaller adult size than Controls despite growing faster on the poor diet ^10,11^. Therefore, we combined adult body size (dry weight) of freshly emerged adults (**Fig S2**) with developmental time data (**Fig 1A**) to estimate mean larval growth rate of each population under both microbiota conditions, following the approach described in ^10^. As expected, we found that inoculation with microbiota increased larval growth rate, but this effect was significantly greater in Control than in Selected populations (**Fig 5A**), suggesting that IIS and/or target of rapamycin (TOR) pathways may respond differently to microbiota (**Fig 5A**).

**Fig 5.**
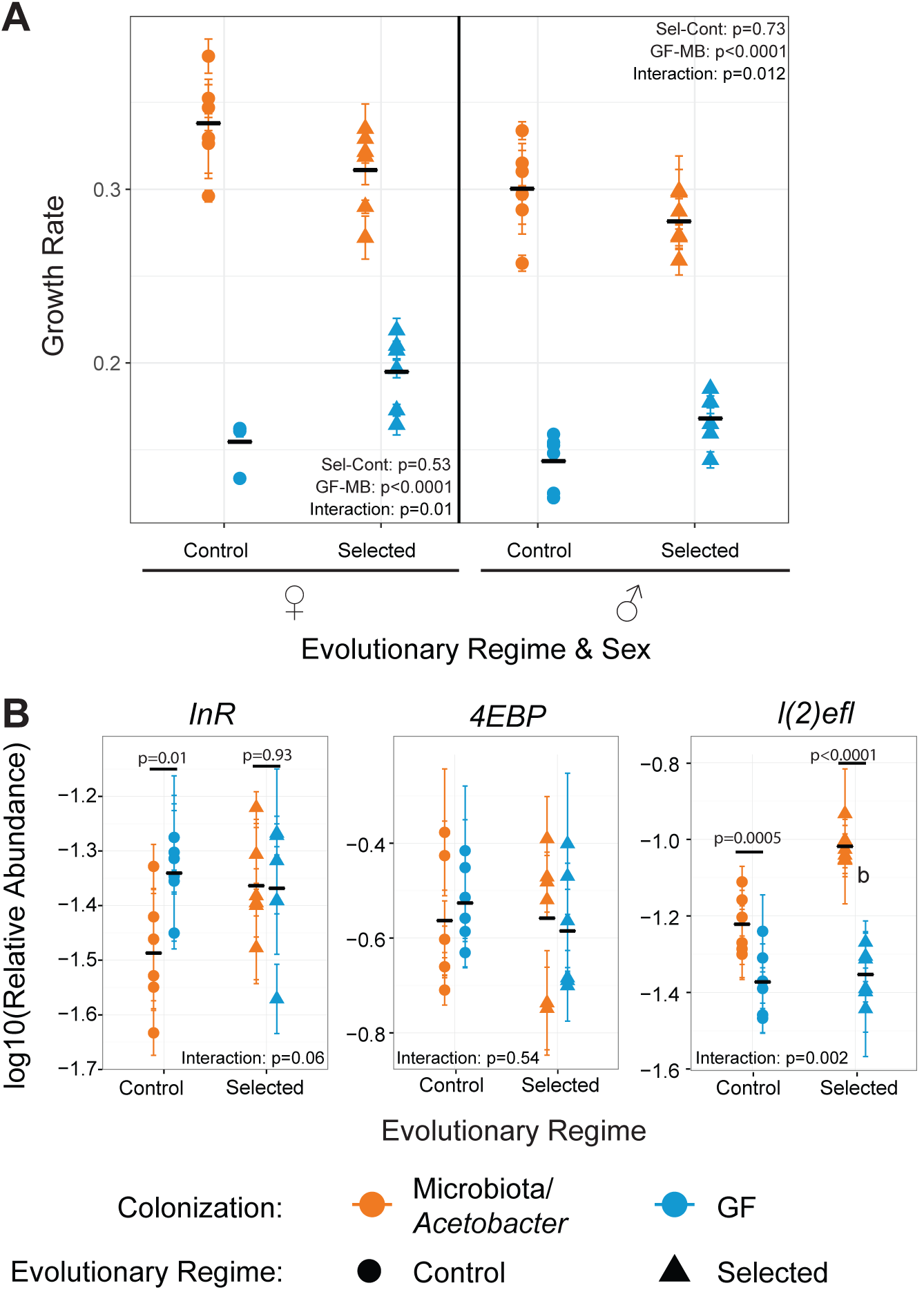
Growth rate and IIS pathway are regulated differently by microbiota association in Selected and Control populations. **A.** Growth rate on poor medium for males and females of Selected and Control populations with and without microbiota. Main effect differences analyzed by GMM are represented in the panel. **B.** Relative abundance (2^-ΔCt^) of different dFOXO targets measured by qRT-PCR from whole Selected and Control larvae that are GF or mono-associated with *Acetobacter* at late 3^rd^ larval instar. Points represent mean±SEM of for 6 populations. Black bars represent the mean for 6 populations. The interaction between colonization and evolutionary regime analyzed by GMM, and significant pairwise contrasts within each regime are represented in the panel. Detailed statistics are presented in Supplementary Table S5.

In *Drosophila*, TOR and IIS pathways control systemic larval growth and dFOXO is the key mediator of IIS in regulating ribosome biogenesis and cellular growth ^12^. dFOXO is a transcription factor that has >900 direct or indirect targets, a part of which respond to nutrient sensing ^16^. To study IIS-dFOXO activity, we analyzed the transcription of three established dFOXO targets, namely *d4EBP, dInR*, and *l(2)efl^17–19^* in late third stage whole larvae upon mono-association with the *Acetobacter* strain isolated from our populations. We observed a significant reduction in *InR* mRNA levels upon colonization by bacteria in Control populations but not in Selected ones (**Fig5B**). Since InR is negatively regulated by insulin-like peptides ^19^, this suggests that Control populations do switch from “low nutrition” to “high nutrition” mode physiologically whereas Selected populations are rather insensitive to inoculation with *Acetobacter* and keep their metabolic state as it is. However, we observed no significant difference in *d4EBP* (**Fig 5B**), indicating that differential dFOXO activity in Selected and Control populations does not occur on all dFOXO targets.

Interestingly, we also saw that the stress response gene *l(2)efl*, known to be involved in lifespan regulation ^17^, was reduced in all populations when they were GF and significantly induced in *Acetobacter* colonized larvae only in Selected populations (**Fig 5B**). This might suggest that Selected populations perceive colonization by *Acetobacter* as a stress signal; however this gene might also play a hitherto unknown role in larval development or nutrition. Taken together these data suggest that dFOXO activates the transcription of a selection of its target differentially in Selected and Control populations in response to colonization by *Acetobacter*.

## Discussion

We set out to study physiological bases of experimental evolutionary adaptation to chronic juvenile malnutrition, expecting that they will involve an improved ability of the animal host to exploit its microbiota. Instead, we found that our experimentally evolved Selected populations became much less dependent on microbiota for their survival and growth in less than 200 generations of evolution on a nutrient-poor larval diet. This is rather surprising, given the well-documented dependence of non-adapted *Drosophila* larvae facing nutrient shortage on benefits provided by gut microbiota ^6–8^. This dependence on microbiota remains strong in our Control populations, which originated from the same base population as Selected populations but do not have a history of laboratory evolution on the poor diet.

It has previously been described in *Drosophila* adults and larvae that *L. plantarum* (mono-associated or as a part of a microbiota community) induces transcription of a set of digestive enzymes ^7^,^20^. Our data from the non-adapted Control populations support the notion that microbiota promote protein digestion and indicate that this effect is not specific to microbiota containing *Lactobacilli* but it also occurs in association with *Acetobacter.* In 2011, Shin et al. showed that benefit conferred by their *Acetobacter* strain arises through the induction of IIS by acetic acid production; yet, adding only acetic acid to the medium does not bring any growth benefit, indicating that other bacterial factors are involved in growth promotion ^8^. Our data complements this view and suggest that enhanced digestion (presumably resulting in improved nutrient acquisition) may be one of the mechanisms mediated by *Acetobacter*, in addition to acetic acid secretion. Interestingly, our data also reveal that even though they are all serine proteases, Trypsins and Jonah proteases respond quite differently to *Acetobacter* colonization, indicating that these two sets may have functional differences.

Enhanced Jonah protease activity has been shown to be sufficient for promoting larval growth upon malnutrition ^7^. This, in combination with our data where GF Selected populations show a higher proteolytic activity mid-3^rd^ larval stage and lower amylase activity throughout the third stage suggests enhancing digestion can be an evolutionary mechanism to insure growth under nutritional stress. In addition to the basal differences that occur at the GF state, we have also shown that microbiota affect digestion differently in Selected and Control populations. In Control populations, we observe a large increase in proteolytic activity as well as a clear decrease in amylase activity upon colonization. This is then accompanied by a significant reduction in InR levels, which is an indication of higher levels insulin-like peptides in the system and thus higher nutrient availability ^19^. In contrast, microbiota have little effect on the protease and amylase activity of Selected populations, whose InR levels appear insensitive to *Acetobacter* colonization. This suggests that Selected populations probably keep their metabolism in a “nutrient shortage” mode in order to continue high nutrient uptake rate. This might also be associated with changes in mitochondrial function and oxidative phosphorylation levels in Selected populations. Previously it was described that larvae that were grown in low (1%) yeast has shown reduced mitochondrial abundance and respiration activity in their fat body ^21^. It remains to be determined if our Selected populations have overcome this defect and if microbiota has a direct influence on mitochondrial abundance and function in evolved and wild type populations.

We found an increase in stress response gene *l(2)efl* transcript levels between control and selected populations upon *Acetobacter* colonization. *l(2)efl* has been described to be upregulated under oxidative stress, heat shock and ionizing radiation and shown to be regulated by a detoxifying ABC-transporter dMRP4 and JNK pathway ^17^,^22^. However the link between stresses induced by larval malnutrition and growth remains to be elucidated.

Together with *InR* data, these colonization and regime specific differences in different genes indicate that dFOXO acts differently in control and selected populations; but since this is not true for translation inhibitor *4EBP* ^23^, differences in upstream signaling mechanisms or different transcription factor partners must be involved. A comprehensive transcriptome analysis would give a more precise picture on the transcriptional changes that occur during adaptation to chronic malnutrition and upon microbiota association.

The genetic basis of natural variation in microbiota-dependent nutritional response was previously studied using *Drosophila* Genetic Reference Panel (DGRP) lines ^24,25^. These studies identified key genes involved in nutritional allocation by microbiota. As expected, genes related to IIS/TOR pathways, as well as JAK-STAT or Notch pathways, were shown to be important for microbiota dependent nutrition ^24^. These genes were identified by genome-wide association studies upon nutritional indices on flies raised on highly rich (10% sugar-yeast diet), which is clearly different than our setup. Despite this, future genomics studies comparing significant SNPs between our Control and Selected populations to the ones identified in these studies will help us understand adaptive forces that shape nutrient acquisition in the presence and absence of microbiota.

Irrespective of its physiological basis, the fact that our Selected populations became much less dependent on microbiota for larval growth and survival under strong nutrient limitations is intriguing from an evolutionary viewpoint. Our quantification of microbiota implies that under their culture regime Selected populations are similarly exposed to microbiota as Control populations, and become colonized by a quantity of bacteria comparable to that resulting from our experimental inoculations. This suggests that the reduced dependence of Selected populations on microbiota is not a consequence of being underexposed to bacteria in the course of their experimental evolution, but a direct effect of adaptation to nutrient shortage under the strong selection imposed by the extremely poor diet. As we have reported ^26^ the Selected populations also evolved a greater susceptibility to the gram-negative intestinal pathogen *Pseudomonas entomophila.* Thus, evolutionary adaptation to nutritional stress may affect interactions between the host and both beneficial and harmful gut microbes.

The relationship between animals and gut microbiota likely goes back hundreds of millions of years, and during this evolutionary time most animals became dependent on gut bacteria for nutritional benefits and various metabolic tasks ^5,27^. In *Drosophila* (and presumably in many other insects with diverse diets) this host-microbiota relationship is less intimate than in mammals or in insects feeding on unbalanced or hard-to-digest diets, such as blood, plant sap or wood ^4^,^5^. Rather than relying on transmission of specialized gut microbes from mother to offspring, *Drosophila* acquire their gut microbiota from the microbial community living on the food substrate ^3^. However, microbiota still exerts its beneficial effect by supplementing food with vitamin B in poor environments and regulating sugar metabolism in high glucose environments ^28^. In addition, *Drosophila* microbiota also stimulates the host immune system, interferes with pathogens, and provides signals to key pathways to which regulate growth and tissue homeostasis ^29^,^30^. And, given that the natural food for *D. melanogaster* is decomposing fruit, the larvae are likely never deprived of those beneficial microbes in nature. It is thus remarkable that the species retained the potential to rapidly evolve a markedly reduced dependence on gut microbiota for fitness under nutritional stress.

## Methods

### Experimentally evolved fly populations and diet

Six replicate Selected and six replicate Control populations were maintained at 20°C and 70% humidity, with 12/12h dark/light cycle on a 21-day generation cycle. Control populations were cultured on standard cornmeal (5%)-yeast (1.25%)-sugar (3% sucrose, 6% glucose) medium and Selected populations were cultured on poor medium containing 1/4 of the nutrients during larval development ^10^. Experimental evolution was carried out as described in detail in ^10^. All 12 populations originated from the same base population. At each generation, eggs were collected on live yeast, leading to contamination of egg surfaces with yeast, which may cause alterations in the gut microbiota of larvae. Eggs were rinsed with tap water to enable egg counting, which dilutes the flies’ natural microbiota and causes environmental contamination. Approximately 200 eggs were collected from adults of each population and distributed on their respective media for larval growth. Upon emergence, adults from all populations were transferred to standard medium supplemented with dry yeast. Experiments were carried out between generation 177 and generation 200. Before each experiment, populations were reared on standard medium for >2 generations to avoid maternal effects. To avoid changes in the conventional recipe, we kept the food clean by boiling it. The food used in the experiments was boiled for >10 min and poured at 78°C in autoclaved fly bottles using tools sterilized with 70% ethanol under the hood.

### Preparation of gnotobiotic larvae

Embryos were collected from an overnight egg laying on orange juice-agar plates supplemented with yeast. Embryos were washed with tap water, sterilized by soaking in 5% bleach for 3 minutes and were rinsed with autoclaved water. 200 eggs were counted on a mesh, under a stereomicroscope, next to a Bunsen burner to avoid further contamination. Counted eggs were transferred to fly bottles containing standard or poor food medium. For the GF treatment, 300 μl of heat inactivated bacteria (developmental time experiment) or sterile PBS (enzymatic activity assays and RT-qPCR experiments) was added on the sterile embryos.

To colonize larvae with microbiota, fecal transplantation was used. Adults (10 males and 10 females) were collected from all populations and kept on standard food for five days. They were transferred on a petri dish with a slice of medium and allowed to defecate for 48 hours. Feces were collected after removal of the medium using an ethanol washed brush in sterile PBS. Feces were filtered through a previously bleached and rinsed mesh and remaining solution was adjusted to a culture turbidity (OD) of 1 to have approximately 10^9^ cells. 300 μl were inoculated on the embryos for colonization.

To mono-associate larvae with *Acetobacter*, bacteria were grown for 48 hours at 30°C under agitation in Man, Ragosa and Sharpe (MRS) medium (Difco, #288110) supplemented with 2.5% D-Mannitol (Sigma, #M1902). Bacteria were harvested by centrifugation at 3000 rpm for 10 minutes and diluted with sterile PBS to reach OD 1. 300 μl of culture was added on sterile embryos.

### Developmental Time and Survival

To measure developmental time and egg-to-pupa survival gnotobiotic animals were prepared as described above. Embryos from 6 Control and 6 selected populations were collected to have 3 biological replicates in each condition (GF vs colonized) and on each food (standard vs poor), resulting in 144 fly bottles to score. Emerging pupae were scored every day to determine larval development time. Three replicate bottles were scored for each population on each condition.

### Adult dry weight and growth rate

The first group of adults emerging from standard or poor food were discarded on the day of emergence and newly emerging ones were collected within 48 hours of eclosion. 10 males and 10 females from each bottle were picked randomly, separated and frozen at -20°C. When the number of adults were not sufficient (valid for GF Control populations on poor food) the procedure was repeated and adults emerged on different days were pooled. If the number of adults was less than 10 the sample was discarded. To determine the dry body weight, flies were dried at 80°C for two days and weighted on a precision balance.

Following ^10^, larval growth rate on poor diet was estimated separately for each sex and population as ln(final size/initial size)/(time available for growth). Final size was the mean dry weight of adults, initial size was assumed to be 0.005 mg, the approximate dry weight of an egg (R. K. Vijendravarma et al, unpublished data). Time available for growth was estimated as the egg-to-adult time minus 48 h to account for time needed for egg hatching, the fact that pupae were scored at 24 h intervals, and the time the larvae spend wandering before pupation (which does not differ between the Selected and Control populations ^31^. While this estimate is necessarily approximate, all conclusions about growth rate were robust to changing the time available for growth by ± 24 h.

### Nucleic acid extraction and qPCR

RNA extractions were performed from three biological replicates of 10 dissected midguts or 10 whole larvae from all six Selected and six Control populations (resulting in 72 gut samples for each time point and 72 whole larval RNA samples) using RNAeasy Mini Kit (Qiagen). Reverse transcription was performed as described in ^7^.

DNA extraction was carried from samples containing 10 surface sterilized (upon washing in sterile water and EtOH) larvae using DNeasy Blood & Tissue Kit (Qiagen) following manufacturer's protocol adapted for insect cells. For conventionally reared lines, larvae were collected from two replicate vials per population. Mono-associated and GF groups were collected from one vial per population.

qPCR was carried out using gene specific primer sets (available as Supplementary information in ^7^ or upon request), using the Power SYBR Green PCR Master Mix (Life technologies, #4368702) under the following conditions: 95°C 10 min, 40 cycles of 95°C, 15 sec and 60°C, 1 min. Melting curve analysis ensured amplification of a single product. Ratios of gene of interest to reference gene (2^-ΔCt^) were log transformed for statistical analysis.

### Protease activity assay

20-50 Whole larvae (equivalent of a volume of 40 μl) were collected from 6 Selected and 6 Control populations in three biological replicates at different time points resulting in 198 individual samples to process. Protease activity was measured using Azocasein assay as described in ^7^, which was optimized for whole larvae.

### Amylase activity assay

Amylase activity was measured using the Amylase Activity Assay Kit (Sigma, #MAK009) following manufacturer's instructions and using the same samples as in the *Protease activity assay.* 50 μl of sample was added to the substrate mix on a 96-well plate. Absorbance at 405 nm was read every 20 min for 17 hours at 25°C. The rate of the reaction, *k* constant, was calculated using non-linear least squares (nls) models in R using function *wrapnls* in package *nlmrt* with the equation: *y=c+A(1-e-^kt^*). The rate was normalized to total protein quantity as for protease activity assay.

### 16S rRNA gene sequencing

Community profiling was from adult feces and poor medium colonizing bacteria during larval stages. Adult feces collection was described in the section “Developmental time and survival” above. Larval medium was washed with 10 ml sterile PBS. The resulting solution was centrifuged for 1 min at 3000 rpm to precipitate the food. The supernatant was re-centrifuged at 13000 rpm for 10 min. Bacterial pellet was resuspended in 1 ml sterile PBS. 5 μl of this suspension was used directly in the PCR to amplify the V1-V2 regions of the 16S rRNA gene, without any DNA extraction. Regions were amplified using the KAPA HiFi HotStart ReadyMix (Kapa Biosystems # KK2601) and primers 8-27F: 5’- TCGTCGGCAGCGTCAGATGTGTATAAGAGACAGAGAGTTTGATCMTGGCTCAG-3' and 339-356R: 5’- GTCTCGTGGGCTCGGAGATGTGTATA-AGAGACAGTGCTGCCTCCCGTAGGAG-3’ including adapter sequences (underlined) for the second PCR round. Three replicate 25 μl PCR reactions containing 10 ng μl^−1^ DNA and 1 μΜ of each primer were carried out under following conditions: 95°C 3 min, 25 cycles of 95°C 30 sec-56°C 15 sec-72°C 30 sec, followed by a final incubation at 72°C for 5 min. Products were pooled from triplicate reactions and verified for amplicon size on a Fragment Analyzer (Advanced Analytical Technologies, Inc.). Libraries were prepared and sequenced at the Lausanne Genome Technology Facilities of the University of Lausanne according to the Illumina 16S Metagenomic Sequencing Library Preparation protocol. Briefly, first round PCR products were cleaned-up using AMPure XB (Beckman Coulter Genomics #A63881) beads. An index PCR was carried out on the purified fraction using a Nextera XT Index Kit (Illumina #FC-131-1001) to produce sequencing libraries. Libraries were again verified by Fragment analyzer, mixed with 20% PhiX library (Illumina #FC-110-3001), and subjected to Illumina MiSeq paired-end sequencing.

All steps of sequence analysis were performed using the QIIME 1.8.0 bioinformatics software ^32^. Raw 300 bp paired-end reads were filtered by size (minimum 100 bp overlap between paired ends) and quality (phred-scores ≥ 20). Chimeric reads were eliminated using the Usearch algorithm ^33,34^ Reads were classified into operational taxonomic units (OTUs) using the open reference OTU clustering pipeline, excluding the pre-filtering step and using the *uclust* method ^33^. Reads were aligned to the Greengenes database ^35^ using PyNAST ^36^ with 99% identity threshold, to have specificity down to the species level. Taxonomies were assigned using the RDP classifier ^37^ and phylogenetic trees were built using FastTree 2.1.3 ^38^.

### Isolation of Acetobacter sp

To isolate *Acetobacter*, media from (randomly chosen) Control #4 and Selected #29 were streaked on MRS-Mannitol plates. A single colony was used to prepare liquid cultures (as described in *Preparation of Gnotobiotic Animals*) and establish glycerol stocks, as well as for 16S rRNA gene full-length amplification using universal primers (sequences available upon request) and KAPA HiFi HotStart ReadyMix. The 16S rRNA gene product was sequenced using Sanger sequencing (GATC Biotech). The obtained sequence was assigned to *Acetobacter* using RDP classifier (https://rdp.cme.msu.edu/classifier/classifier.jsp). To make sure that we isolated the dominant strain, which was detected during community profiling, we aligned sequences using APE Software.

### Statistical Analysis

Univariate analysis was performed using general linear mixed models (GMM) using Satterthwaite approximation for the degrees of freedom (Proc Mixed of SAS v. 9.3). Multivariate analysis was done using “ade4” package in R ^39^. Evolutionary regime (Selected or Control) and microbiota treatment (germ-free or colonized) were fixed factors; time point was also a fixed factor except for enzyme activity assays, where more than two time points were included. Replicate populations were treated as a random factor nested in evolutionary regimes. A priori pairwise contrasts were performed within the framework of the GMM (using the Slices option of Proc Mixed). Detailed output of all analyses can be found in **Supplementary Tables S1-S5**.

## Author Contributions

BE and TJK designed the experiments. BE and SK performed the experiments. BE and TJK analyzed the data. BE, JvdM and TJK wrote the article.

## Acknowledgements

This work was supported by an interdisciplinary grant from Faculty of Biology and Medicine, University of Lausanne, to TJK and JRvdM, and by a Swiss National Science Foundation grant to TJK.

## Competing and financial interests

We declare no competing or financial interests.

